# Three cytochrome P450 from *Nicotiana attenuata* play key roles in triterpene biosynthesis

**DOI:** 10.1101/2024.05.10.593601

**Authors:** Caiqiong Yang, Rayko Halitschke, Sarah E. O’Connor, Ian T. Baldwin

## Abstract

Pentacyclic triterpenoids, recognized for their natural bioactivity, display complex spatiotemporal accumulation patterns within the ecological model plant, *Nicotiana attenuata*. Despite their ecological significance, the underlying biosynthetic enzymes and functional attributes of triterpenoid synthesis in *N. attenuata* remain unexplored. Three multifunctional cytochrome P450 monooxygenases (NaCYP716A419, NaCYP716C87, NaCYP716E107) from *N. attenuata* were shown to oxidize the pentacyclic triterpene skeleton as evidenced by heterologous expression in *Nicotiana benthamiana*. NaCYP716A419 catalyzed a consecutive three-step oxidation reaction at the C28 position of β-amyrin/lupeol/lupanediol, yielding the corresponding alcohol, aldehyde, and carboxylic acid. NaCYP716C87 hydroxylated the C2α position of β-amyrin/lupeol/lupanediol/erythrodiol/oleanolic acid/betulinic acid, while NaCYP716E107 hydroxylated the C6β position of β-amyrin/oleanolic acid. Three CYP716 enzymes are highly expressed in flowers and respond to induction by ABA, MeJA, SA, GA_3_, and abiotic stress treatments. Using VIGS technology, we revealed that silencing of NaCYP716A419 affects the growth and reproduction of *N. attenuata*, suggesting the ecological significance of these specialized metabolite biosynthetic steps.

**One-sentence summary:** Three CYP716 enzymes diversify *N. attenuata’s* triterpenoid sector with potential roles in growth and development.

## Introduction

Triterpenoids, a class of isoprenoid compounds, are universally present in all eukaryotic organisms. They constitute a highly diverse group of polycyclic molecules with a wide range of biological functions, spanning both primary to secondary metabolism. These compounds are elaborated not only in medicinal plants, like licorice and ginseng (Dou et al., 2001; Tran et al., 2001; Seki et al., 2008; Seki et al., 2011), but are also commonly found in crops such as legumes and oats (Geisler et al., 2013; Hu et al., 2021; Yu et al., 2022). Triterpenoids often serve as integral components of plant defense mechanisms, playing central roles in direct defense responses (González-Coloma et al., 2011; Leveau et al., 2019). Their frequent inclusion in traditional medicinal practices underscores their long-recognized pharmacological significance (Wang, 2021; Reed et al., 2023). Furthermore, triterpene saponins are extensively used in various food, cosmetics, and pharmaceutical industrial sectors (Güçlü-Üstündağ and Mazza, 2007; Sharma et al., 2023). Therefore, elucidation of the potential pathways for triterpenoid biosynthesis at a molecular level can facilitate the engineering of these pathways into heterologous hosts or enhance their production in native hosts to improve crop resilience.

Triterpene biosynthesis initiates with the acetylation of Coenzyme A (Co-A), proceeding through the mevalonic acid pathway with various oxidative cyclizations, catalyzed by oxidosqualene cyclases (OSCs), producing a variety of triterpene structures. Currently identified OSCs include β-amyrin and lupeol synthase (Khakimov et al., 2015; Moses et al., 2015; Jo et al., 2017), with some multifunctional OSCs reported to synthesize unique triterpene scaffolds, such as lupanediol, taraxasterol, δ-amyrin, germanicol, and others (Ito et al., 2011; Wang et al., 2011; Moses et al., 2015; Andre et al., 2016; Srisawat et al., 2019). These triterpene scaffolds are subsequently decorated through modifications, including additions of hydroxyl, ketone, aldehyde, or carboxyl groups, and glycosylations.

Cytochrome P450 monooxygenases (CYPs) perform several modifications of triterpene scaffolds which can occur at various positions (Ghosh, 2017; Romsuk et al., 2022). To date, approximately fifty P450 enzymes have been reported to act on plant pentacyclic triterpene scaffolds, the majority belong to the CYP716 family. Additionally, some members of the CYP51, CYP71, CYP72, CYP87, CYP88, and CYP93 families have also been reported to modify pentacyclic triterpenes (Ghosh, 2017). The biochemical properties of multiple members of the CYP716 family have been characterized (Carelli et al., 2011; Moses et al., 2015; Miettinen et al., 2017; Misra et al., 2017; Yasumoto et al., 2017; Sandeep et al., 2019; Romsuk et al., 2022). Most catalyze a series of three consecutive oxidation reactions at the C28 position of β-amyrin/lupeol scaffolds, resulting in the sequential introductions of hydroxyl, aldehyde, and carboxyl groups at the C28 position (Carelli et al., 2011; Fukushima et al., 2011; Misra et al., 2017). Furthermore, certain plant CYP716A genes also catalyze oxidation reactions at carbon atoms other than C28 in β-amyrin scaffolds. For instance, in *Arabidopsis thaliana*, AtCYP716A2 catalyzes hydroxylation at C22α (Yasumoto et al., 2016), while in *Artemisia annua*, AaCYP716A14v2 catalyzes oxidations at C3 (Moses et al., 2015). AcCYP716A111 from *Aquilegia coerulea*, and PgCYP716A141 from *Platycodon grandiflorus* both catalyze hydroxylations at C16β (Miettinen et al., 2017). Members of the CYP716E subfamily, such as SlCYP716E26 from *Solanum lycopersicum* and CaCYP716E41 from *Centella asiatica*, have been biochemically characterized as C6β-hydroxylases that accept α/β-amyrin and oleanolic/ursolic/maslinic acids as substrates (Miettinen et al., 2017; Yasumoto et al., 2017). CaCYP716C11 from *C. asiatica* and OeCYP716C67 from *Olea europaea* catalyze C2α hydroxylations of oleanolic acid, 6β-hydroxy-oleanolic acid, or ursolic acid (Miettinen et al., 2017; Alagna et al., 2023). Furthermore, GuCYP88D6 from licorice catalyzes C11 oxidations (Seki et al., 2008), while GmCYP93E3 catalyzes C24 hydroxylations of β-amyrin (Seki et al., 2008; Moses et al., 2014). In monocotyledonous plants such as oats, AsCYP51H10 is a multifunctional enzyme, catalyzing both hydroxylations and epoxidations of β-amyrin, resulting in the formation of 12,13β-epoxy-3β,16β-dihydroxy-oleanane (Geisler et al., 2013). Although the biosynthetic pathways of triterpenes have been elucidated in some plants, many of the enzymes responsible for triterpene structural modifications remain to be fully elucidated in most plants.

*Nicotiana attenuata* is a native annual wild tobacco species that grows in the Great Basin Desert, Utah, USA, and is a model for native plant-environment ecological interactions (Kessler and Baldwin, 2004; Joo et al., 2021; Yang et al., 2023; You et al., 2023). While triterpenoid compounds have been reported from *Nicotiana* species (Popova et al., 2018; Popova et al., 2019; Popova et al., 2020), our understanding of their biosynthesis remains rudimentary. Previously, we reported that *N. attenuata* inducibly accumulates pentacyclic triterpenoid compounds, primarily derived from lupane and oleanane scaffolds, in young plant organs or flowers. We identified NaOSC1 as being required for the biosynthesis of the triterpene scaffolds, lupeol, β-amyrin, lupanediol, dammarenediol II, and taraxasterol, while NaOSC2 predominantly synthesizes β-amyrin (Yang et al., 2023). Here, we identify and characterize *N. attenuata* cytochrome P450 enzymes from 6 distinct subfamilies: CYP716A (NaCYP716A419, NaCYP716A420), CYP716E (NaCYP716E107, NaCYP716E108), CYP716D (NaCYP716D93, NaCYP716D94), CYP716C (NaCYP716C87), CYP716H (NaCYP716H6), and CYP88B (NaCYP88B4). Through heterologous expression in *N. benthamiana*, we found NaCYP716A419, NaCYP716C87, and NaCYP716E107 to be triterpene modifying enzymes, with NaCYP716A419 catalyzing a consecutive three-step oxidation reaction at the C28 position of β-amyrin, lupeol, and lupanediol; NaCYP716C87 hydroxylating the C2α position of β-amyrin, erythrodiol, oleanolic acid, lupeol, betulinic acid, and lupanediol; and NaCYP716E107 hydroxylating the C6β position of β-amyrin or oleanolic acid. Additionally, using Tobacco Rattle Virus (TRV)-induced gene silencing (VIGS), we further characterized that NaCYP716A419, NaCYP716E107, and NaCYP716C87 are potentially involved in *N. attenuata’s* growth and development.

## Results

### *N. benthamiana*-based *in vivo* screening of candidate P450 enzymes identifies three functional enzymes

Based on the cytochrome P450 hidden Markov model, PF00067, we identified a total of 234 complete P450 cytochrome enzymes in *N. attenuata*, among which, ten were annotated as β-amyrin oxidases (**Fig. S1**). Microarray data sourced from the *Nicotiana attenuata* Data Hub (http://nadh.ice.mpg.de/NaDH/) revealed that *NIATv7_g15098* and *NIATv7_g07976* were predominantly expressed in roots, *NIATv7_g01943* was primarily expressed in various floral organs, and *NIATv7_g06757* and *NIATv7_g33874* were mainly expressed in seeds, with some expression in floral organs. In contrast, *NIATv7_g18201* and *NIATv7_g15096* were expressed in both roots and floral organs, while *NIATv7_g17429*, *NIATv7_g33423*, and *NIATv7_g03976* were primarily expressed in stems and leaves (**Fig. 1A**). We conducted a phylogenetic analysis of these candidate P450 enzymes in comparison to known P450 oxidases involved in triterpenoid biosynthesis (**Supplementary Table S1**). The results revealed that NIATv7_g01943 and NIATv7_g18201 clustered on the same branch as MtCYP716A12, known for C28 oxidation activity in *Medicago truncatula* (**Fig. 1A**)*. NIATv7_g01943* and *NIATv7_g18201* share 87 and 85% amino acid sequence similarities with CYP716A12, respectively (**Fig. S2**). NIATv7_g15096 and NIATv7_g15098 clustered with SlCYP716E26, which exhibits C6β oxidation activity (**Fig. 1A**). Their amino acid sequences were 62 and 63% identical to SlCYP716E26, with similarities of 80 and 81% (**Fig. S2**). NIATv7_g06757 and NIATv7_g33874 clustered with GuCYP88D6 (**Fig. 1A**), and their amino acid sequences were 40 % identical to GuCYP88D6, with similarities of 61% (**Fig. S2**). NIATv7_g07976 clustered on the same branch with CaCYP716C11 (**Fig. 1A**) with 63% identity and 78% similarity (**Fig. S2**).

**Fig. 1.**
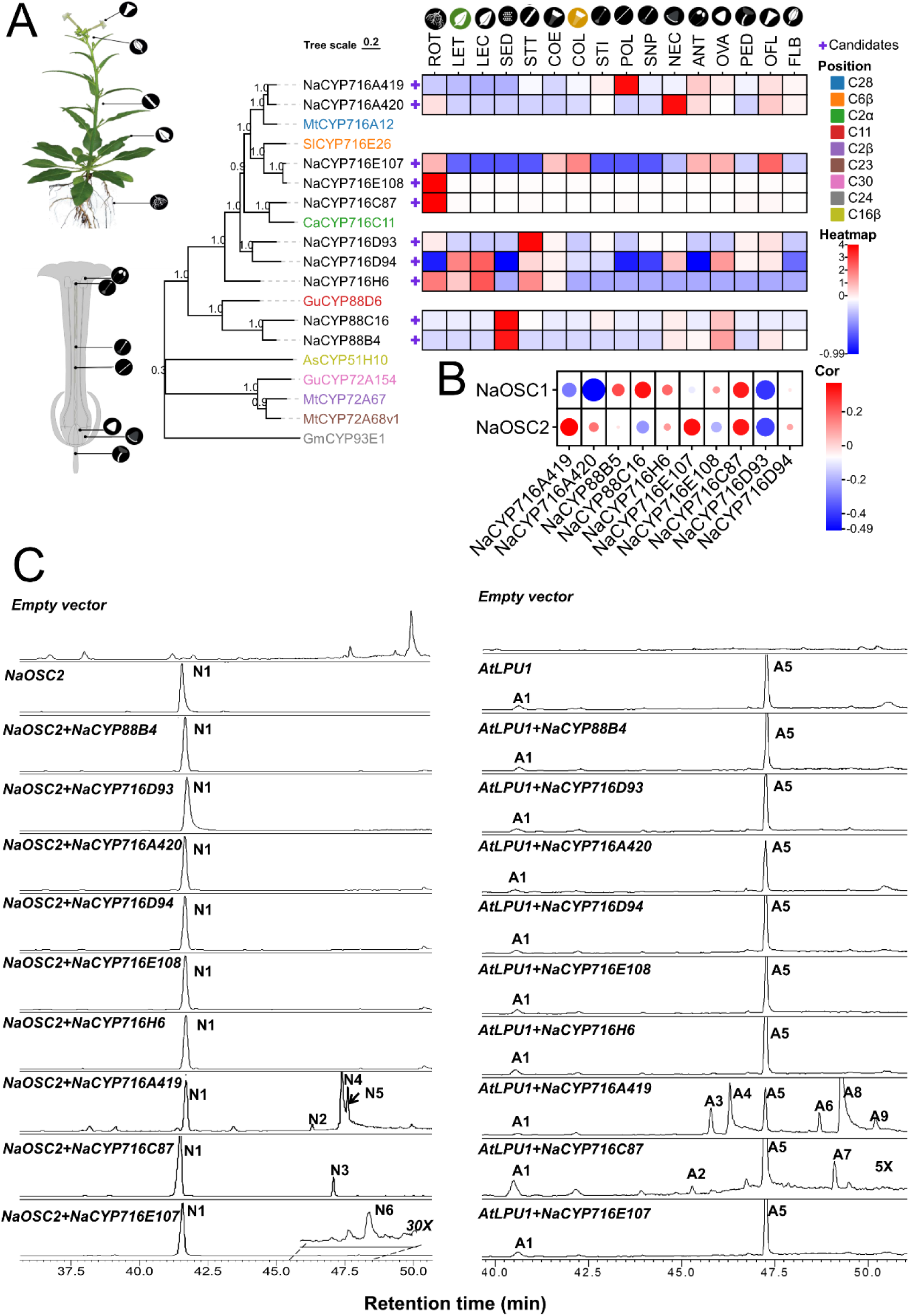
Screening of cytochrome P450 enzymes from the triterpenoid biosynthesis pathway in *N. attenuata*. **(A)** Phylogenetic tree and tissue-specific expression of candidate cytochrome P450 enzymes. The protein sequences of known triterpenoid biosynthetic enzymes were obtained from the NCBI Genebank, while the protein sequences and microarray data of candidate enzymes were sourced from the *Nicotiana attenata* Data Hub (Brockmöller et al., 2017). ROT: root treated with *Manduca sexta* oral secretion, LET: leaf treated with *M. sexta* oral secretion, LEC: leaf control, SED: seed; STT: stem treated with *M. sexta* oral secretion, COE: corolla early, COL: corolla late, STI: stigma, POL: pollen tubes, SNP: style without pollination, ANT: Anthers, NEC: nectaries, OVA: ovary, PED: pedicels, OFL: opening flower, FLB flower buds. In the phylogenetic tree, the color-coded P450 enzymes represent those already reported to have triterpenoid modification functions. **(B)** Correlation analysis of candidate genes with oxidosqualene cyclase tissue expression data. The microarray data used for correlation analysis was obtained from the *N. attenuta* Data Hub. The depth of color represents the correlation coefficient, while the data points’ size indicates the significance level. **(C)** GC-MS chromatograms of extracts of *N. benthamiana* leaves co-expressing CYP450 candidate genes and oxidosqualene cyclase. “30x” and “5x” respectively indicate that the chromatograms had been magnified by a factor of 30x and 5x, respectively. Triterpene compound obtained by heterologous expression are labeled as peaks N1-6 (NaOSC2+CYP) and A1-9 (AtLPU1+CYP). Each treatment was conducted with at least three replications.

The remaining three genes, NIATv7_g03976, NIATv7_g33423, and NIATv7_g17429, displayed low sequence similarities of less than 40% when compared to known triterpene oxidases. These ten candidate CYP450 enzymes were submitted to the P450 Nomenclature Committee for naming and NIATv7_g01943 and NIATv7_g18201 were designated as NaCYP716A419 and NaCYP716A420, in the CYP716A subfamily. NIATv7_g15096 and NIATv7_g15098 were designated as NaCYP716E107 and NaCYP716E108, belonging to the CYP716E subfamily. NIATv7_g07976 was designated NaCYP716C87, a member of the CYP716C subfamily. NIATv7_g03976 was designated as NaCYP716H6 in the CYP716H subfamily. NIATv7_g33423 and NIATv7_g17429 were placed in the CYP716D subfamily and designated as NaCYP716D93 and NaCYP716D94, respectively. Lastly, NIATv7_g06757 and NIATv7_g33874 were placed in the CYP88C and CYP88B subfamilies and named NaCYP88C16 and NaCYP88B4, respectively (**Supplementary Table S2**).

Previously, our research identified the roles of NaOSC1 and NaOSC2 in the biosynthesis of triterpene scaffolds in *N. attenuata* (Yang et al., 2023). We conducted Spearman’s correlation analysis to examine associations among the expressions of *Naosc1* and *Naosc2* in different tissues and CYP450 candidates. Strong positive correlations were found between *Nacyp716a419*, *Nacyp716e107*, and *Nacyp716c87* with *Naosc2*, and between *Nacyp716c87* and *Nacyp88c16* with *Naosc1* (**Fig. 1B**). From these correlations, we hypothesized that these CYP450 enzymes might function in the down stream pathway of triterpenes after NaOSC1 or NaOSC2, respectively. Except for NaCYP88C16, all were successfully cloned into the heterologous expression vector 3Ω1 (Cárdenas et al., 2019; Hong et al., 2022).

To determine the oxidation activity of these P450 enzymes towards simple triterpenes in *N. attenuata*, we conducted combination experiments in which these candidate genes were co-expressed with OSCs. In *N. benthamiana*, heterologous expression of NaOSC2 results in abundant production of β-amyrin, while heterologous expression of NaOSC1 yields a diverse array of products, including 11 compounds represented by lupeol and lupanediol. Due to the diverse and generally low levels of products obtained from the heterologous expression of NaOSC1 in *N. benthamiana* (Yang et al., 2023), we opted to substitute NaOSC1 with an enzyme from *Arabidopsis thaliana*, AtLPU1, known to produce higher levels of lupeol and lupanediol when heterologously expressed in *N. benthamiana* (Segura et al., 2000). Leaves expressing NaOSC2 or AtLUP1 were used as negative controls. GC-MS analysis revealed distinct chromatographic peaks for NaCYP716C87, NaCYP716A419, and NaCYP716E107 compared to the control group (**Fig.1C, Fig.S3**). Specifically, the expression of NaCYP716C87 and NaCYP716A419 generated novel peaks with high yields when co-expressed with NaOSC2 (peaks N2-5) or AtLUP1 (peaks A2-4 and A6-9), whereas NaCYP716E107 showed only weak activity when co-expressed with NaOSC2 (peak N6) but not with AtLUP1 (**Fig. 1C, Fig.S3**).

### NaCYP716A419 is a C28 oxidase

Comparisons of retention times and Electron Impact-Mass Spectrometry (EI-MS) spectra with authentic commercial standards, revealed that the EI-MS and retention times of peaks N2 and N4 matched those of erythrodiol and oleanolic acid (**Fig. 2A, Supplementary Table S3**). Additionally, the EI-MS and retention times of peaks A3 and A4 matched those of betulin and betulinic acid (**Fig. 2B, Supplementary Table S4**). Due to the absence of reliable standards, our identification of oleanolic aldehyde (N5) is based on existing EI-MS data and the relative elution order of these triterpenes in gas chromatography (Misra et al., 2017) **(Supplementary Table S3)**. In *N. benthamiana* leaves co-expressing NaOSC2 and NaCYP716A419, three new peaks were observed (**Fig. 2A**), corresponding to three C28 oxidized products of β-amyrin (N1), namely erythrodiol (N2), oleanolic aldehyde (N5), and oleanolic acid (N4). Meanwhile, we also compared the triterpenoid profiles when NaCYP716A419 was co-expressed with either EV or NaOSC2 in *N. benthamiana* leaves, demonstrating that the β-amyrin substrate present in *N. benthamiana* leaf tissue does not yield detectable levels of erythrodiol (N2), oleanolic aldehyde (N5), and oleanolic acid (N4) (**Fig. S4A**). Expression of the empty vector in *N. benthamiana* leaves and supplemented with erythrodiol (N2) as a substrate also did not result in detectable levels of oleanolic aldehyde (N5) and oleanolic acid (N4), indicating that oleanolic aldehyde (N5) and oleanolic acid (N4) are not products of endogenous enzymes in *N. benthamiana* leaves (**Fig. S4B**).

**Fig. 2.**
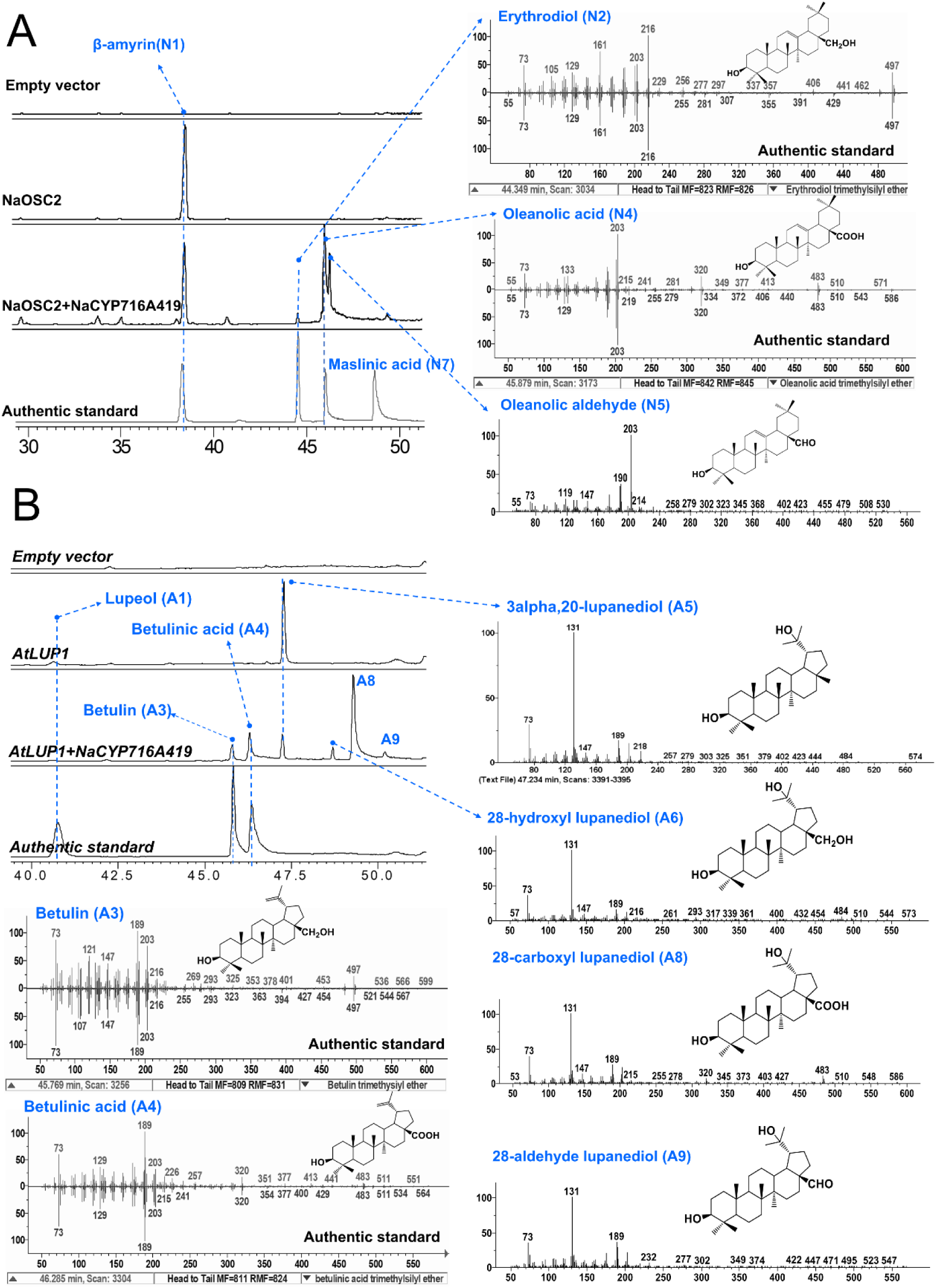
Characterization of CYP716A419 as a C28 oxidase. **(A)** Gas chromatography (GC)-MS profiles (with Selective Ion Monitoring (SIM) at m/z: 189, 203, 216, 320) and electron impact-mass spectrometry (EI-MS) spectra of trimethylsilylated triterpenoids of extracts of *N. benthamiana* leaves co-expressing CYP716A419 with NaOSC2. **(B)** GC-SIM-MS profiles and electron impact-mass spectrometry (EI-MS) spectra of trimethylsilylated triterpenoids of extracts of *N. benthamiana* leaves co-expressing CYP716A419 with AtLUP1. Each treatment was conducted with at least three replications.

In *N. benthamiana* leaves co-expressing AtLUP1 and NaCYP716A419, five new peaks were observed (**Fig. 2 B**), representing two C28 oxidized forms of lupeol (A1), betulin (A3), and betulinic acid (A4), as well as three putative C28 oxidized forms of lupanediol (A5), including 28-hydroxy lupanediol (A6:(1R,3aS,5aR,5bR,9S,11aR)-3a-(hydroxymethyl)-1-(2-hydroxypropan-2-yl)-5a,5b,8,8,11a-pentamethylicosahydro-1H-cyclopenta[a]chrysen-9-ol), 28-adehyde lupanediol (A9: (1R,3aS,5aR,5bR,9S,11aR)-9-hydroxy-1-(2-hydroxypropan-2-yl)-5a,5b,8,8,11a-pentamethylicosahydro-3aH-cyclopenta[a]chrysene-3a-carbaldehyde), and 28-carboxy lupanediol (A8: (1R,3aS,5aR,5bR,9S,11aR)-9-hydroxy-1-(2-hydroxypropan-2-yl)-5a,5b,8,8,11a-pentamethylicosahydro-3aH-cyclopenta[a]chrysene-3a-carboxylic acid). These identifications were based on characteristic ions in the EI-MS spectra and the oxidative function of NaCYP716A419 (**Supplementary Table S4**). From these data, we infer that NaCYP716A419 is an enzyme capable of catalyzing oxidation reactions at the C28 position of β-amyrin (N1), lupeol (A1), and lupanediol (A5) triterpene scaffolds, resulting in the formation of hydroxyl, aldehyde, and carboxyl groups (**Fig. 2A, B**)

### NaCYP716C87 is a C2α hydroxylase

The phylogenetic analysis (**Fig. 1**) revealed that NaCYP716C87 clustered with CaCYP716C11, a cytochrome P450 monooxygenase from *Centella asiatica* reported having C2α oxidation activity of oleanolic acid to produce the product, maslinic acid (Miettinen et al., 2017). The high degree of amino acid sequence identity (63%) and similarity (78%), suggested a similar function. To verify the C2α oxidation activity of NaCYP716C87, we heterologously expressed the NaCYP716C87 enzyme in *N. benthamiana* leaves and fed them with oleanolic acid substrate. The results revealed peak N7, with retention time and GCMS spectrum consistent with the maslinic acid standard (**Fig. 3A**). When NaOSC2, NaCYP716A419, and NaCYP716C87 are simultaneously expressed in *N. benthamiana*, a strong maslinic acid (N7) peak was detected as well as a notable decrease in oleanolic acid compared to leaves expressing only NaOSC2 and NaCYP716A419 (**Fig. 3B, Fig. S5**). This indicates that NaCYP716C87 also functions as a C2α hydroxylase. Concurrently, co-expression of NaOSC2, known to possess β-amyrin synthase activity, with NaCYP716C87 in *N. benthamiana* leaves resulted in the production of β-amyrin and another new peak, N3 (**Fig. 1B and Fig. 3C**). The derivatization pathway of β-amyrin standard β-amyrin-TMS, after undergoing EI-MS fragmentation, is illustrated in **Fig. S7**. β-amyrin-TMS undergoes retro-Diels–Alder cleavage (RDA), resulting in m/z 218 and m/z 279. m/z 218 looses a methyl group and gains an electron to yield m/z 203, while m/z 279 looses a TMSOH and forms a double bond to yield m/z 189. When the C2α position is substituted with a hydroxyl group, as in maslinic acid (**Fig. S11**), its trimethylsilylated derivative undergoes RDA to yield m/z 320 and m/z 367. m/z 320 looses TMSCOOH and gains a proton to yield m/z 203, while m/z 367 looses a TMSOH and forms a double bond to yield m/z 277. Furthermore, m/z 277 continues to loose three methyl groups and gain three protons to yield m/z 235. N3 exhibits m/z 218, 203, and 189, which is similar to the fragmentation pattern of β-amyrin, and similarily, m/z 277 and 235 match the steps of AB ring loss ions after RDA cleavage of maslinic acid containing a hydroxyl group at the C2α position. By comparing their EI-MS spectra and fragmentation pathways with those of β-amyrin and maslinic acid (**Fig. S7, S11, and Fig. 3D**), N3 was inferred to be 2α-hydroxyl-β-amyrin. Furthermore, feeding erythrodiol to *N. benthamiana* leaves expressing NaCYP716C87 led to the appearance of another new peak, N8 (**Fig. 3C**). By comparing its EI-MS spectra with those of maslinic acid and erythrodiol (**Fig. S8, Fig. S11, and Fig.3 D**), N8 was suggested to be 2α-hydroxyl-erythrodiol.

**Fig. 3.**
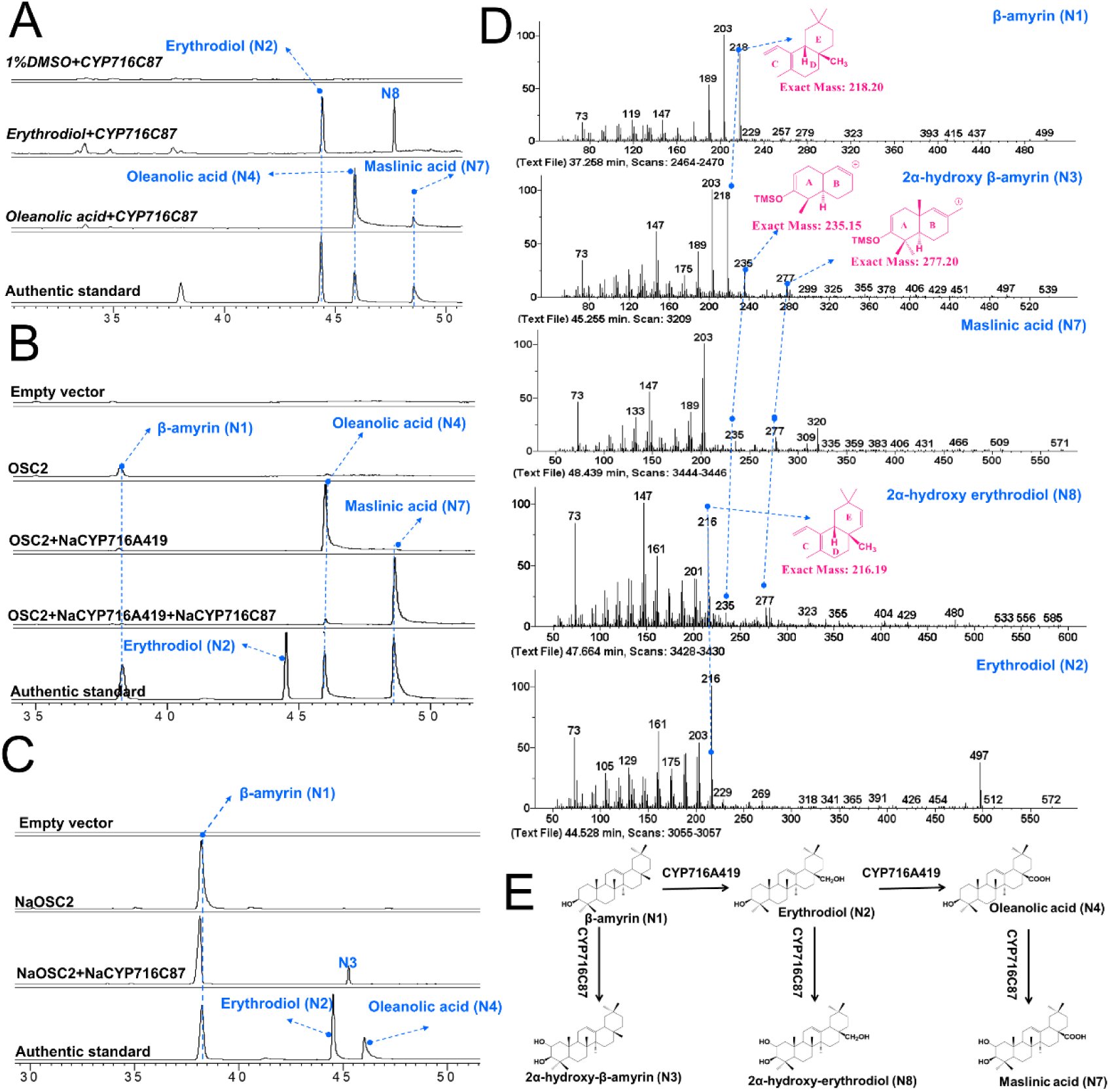
CYP716C87 is a C2α oxidase. **(A)** GC-SIM MS (m/z: 189, 203, 147, 320) chromatograms of extracts from *N. benthamiana* leaves expressing only CYP716C87 and supplemented with erythrodiol (N2) and oleanolic acid (N4) substrates (100 μM in 1% DMSO). 1% DMSO was added to *N. benthamiana* leaves expressing CYP716C87 as a control. **(B)** GC-SIM MS (m/z: 189, 203, 147, 320) chromatograms of extracts of *N. benthamiana* leaves co-expressing NaOSC2, NaCYP716A419, and NaCYP716C87. Leaves expressing empty vectors, NaOSC2 and NaOSC2+ NaCYP716A419 were employed as controls. **(C)** GC-SIM MS (m/z: 189, 203, 147, 320) chromatograms of extracts of *N. benthamiana* leaves co-expressing NaOSC2 and NaCYP716C87. Leaves expressing empty vectors or only NaOSC2 were employed as controls. **(D)** Electron impact-mass spectrometry (EI-MS) spectra of trimethylsilylated 2α-hydroxy-β-amyrin (N3), maslinic acid (N7), 2α-hydroxy-erythrodiol (N8), β-amyrin(N1), and erythrodiol. **(E)** Structures of the substrate, intermediates, and products of the oxidation reaction step from β-amyrin to maslinic acid by CYP716A419 and CYP716C87. Each treatment was conducted with at least three replications.

In *N. benthamiana* leaves expressing AtLUP1, three distinct peaks, namely N1, A1, and A5, were observed (**Fig.4**). Peaks A1 and A5 correspond to the major products of AtLUP1, lupeol, and lupanediol, respectively (Segura et al., 2000). Co-expression of AtLUP1 and NaCYP716C87 in *N. benthamiana* resulted in the appearance of two new peaks, designated as A2 and A7 (**Fig. 4, Fig. S3**). Comparison of their EI-MS spectra and fragmentation pathways with those of lupeol (A1) (**Fig. S12**), lupanediol (**Fig.S13**), and maslinic acid (**Fig.S11**) revealed characteristic fragments indicative of potential C2α hydroxylation. Additionally, peak A2 exhibited the characteristic fragment 189 of lupeol, while peak A7 displayed the characteristic fragment 131 of lupanediol (**Fig. 4**, **Fig. S12-S13**). Therefore, we inferred that peak A2 corresponds to 2α-hydroxy-lupeol and peak A7 corresponds to 2α-hydroxy lupanediol. Upon co-expression of AtLUP1, NaCYP716C87, and NaCYP716A419 in *N. benthamiana*, peaks A2 and A7 disappeared, while peaks A10 and A11 emerged (**Fig. 4, Fig. S6**). Based on the EI-MS spectra and inferred fragmentation pathways of oleanolic acid (N4, **Fig. S10**), betulinic acid (A4, **Fig. S15**), and maslinic acid (N7, **Fig. S11**), compounds bearing a C28 carboxyl group exhibit a characteristic fragment at m/z 320. Combining the previous findings (**Fig. 3**), peak A10 is inferred to be 2α-hydroxyl-betulinic acid (alphtolic acid), whereas peak A11 is suggested to be 2α-hydroxy-28-carboxyl lupanediol ((1R,3aS,5aR,5bR,9R,11aR)-9,10-dihydroxy-1-(2-hydroxypropan-2-yl)-5a,5b,8,8,11a-pentamethylicosahydro-3aH-cyclopenta[a]chrysene-3a-carboxylic acid).

**Fig. 4.**
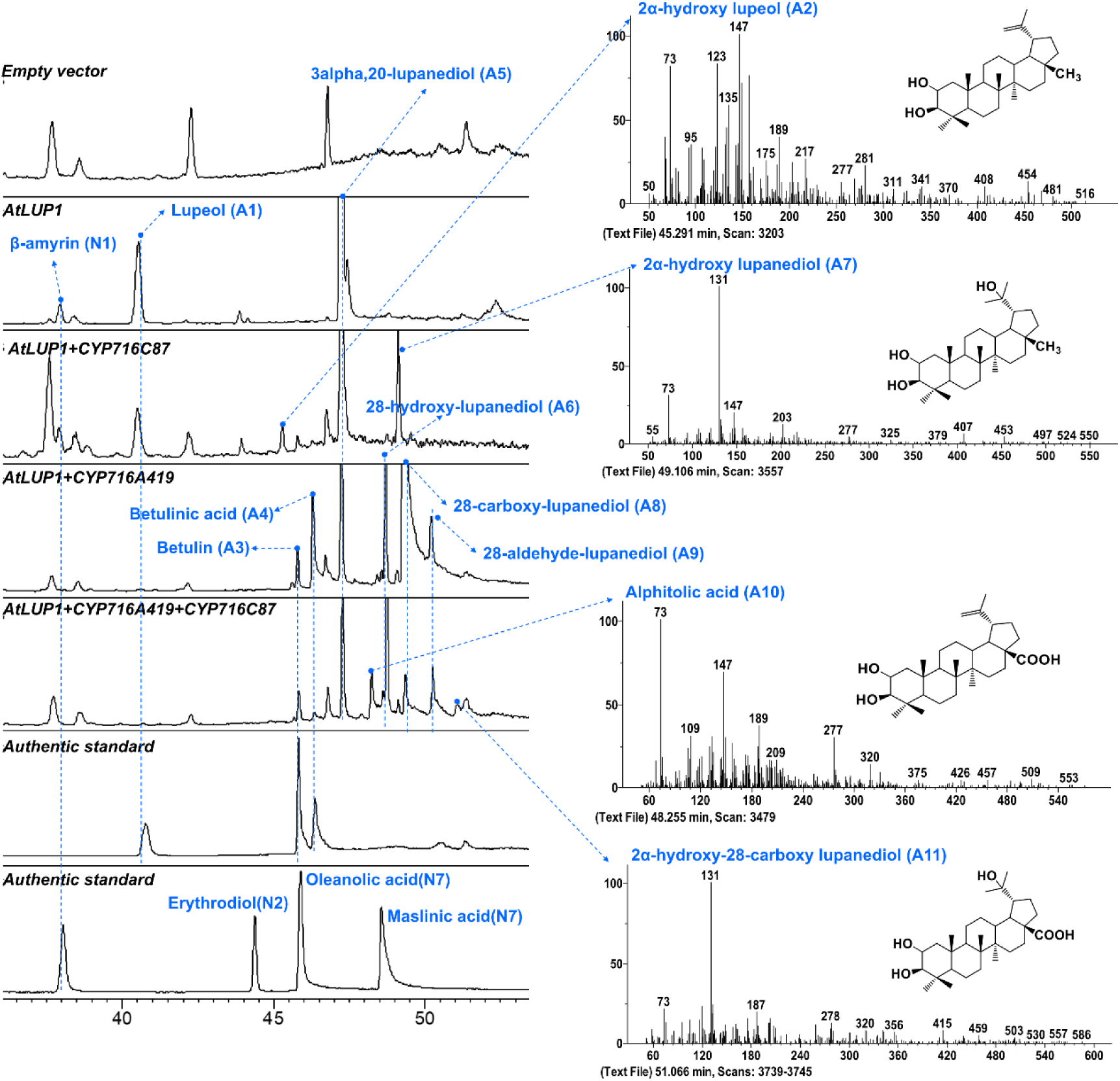
GC-MS chromatograms and EI-MS spectra of TMS-derivitization of extracts of *N. benthamiana* leaves co-expressing AtLUP1 and NaCYP716C87. Leaves expressing empty vectors, AtLUP1, and AtLUP1+ NaCYP716A419 were employed as controls. Each treatment was conducted with at least three replications.

In summary, CYP716C87 functions as a C2α hydroxylase, which accepts β-amyrin (N1), erythrodiol(N2), oleanolic acid (N4) (**Fig.3E**), as well as lupeol (A1), lupanediol (A5), betulinic acid (A4), and 28-carboxy lupanediol (A8) as substrates (**Fig. 4**).

### NaCYP716E107 is a C6β hydroxylase

Amino acid sequence analysis revealed a 62% identity and 80% similarity between NaCYP716E107 and SlCYP716E26 (**Fig. S16**), a hydroxylase that catalyzes the β-amyrin C6β hydroxylation reaction and produces daturadiol as a product (Yasumoto et al., 2017).To determine if NaCYP716E107 has C6β hydroxylation functionality given the lack of daturadiol standards, we cloned SlCYP716E26 from tomato roots. Then, we conducted heterologous expression in *N. benthamiana* by co-expressing the enzyme individually with NaOSC2 or in combination: NaOSC2/NaCYP716A419 (**Fig. 5A, B**). The leaves expressing NaOSC2 and SlCYP716E26 produced two peaks, β-amyrin (N1) and N6 (**Fig. 5A, Fig. S17**). By comparing the retention times and EI-MS spectra with those reported products for SlCYP716E26 (Yasumoto et al., 2017), we confirmed that N6 is daturadiol (**Supplementary Table S3**). Leaves co-expressing NaOSC2 and NaCYP716E107 yielded a peak at the same position. Upon GC-MS analysis, the peak matched with daturadiol (N6) obtained from the co-expression of NaOSC2 and SlCYP716E26. However, the yield was reduced by 95% compared to when SlCYP716E26 was co-expressed with NaOSC2 (**Fig. 5A and Fig. S17A**). CaCYP716E41 from *Centella asiatica* is another enzyme demonstrated to catalyze C6β hydroxylation of triterpenes. It shares high sequence similarity with NaCYP716E107 and SlCYP716E26 (**Fig. S16**). However, unlike NaCYP716E107 and SlCYP716E26, CaCYP716E41 does not catalyze C6β hydroxylation of β-amyrin. Instead, it catalyzes oleanolic acid to yield two peaks: 6β-hydroxyl-oleanolic acid and putative incompletly derivatized 6β-hydroxyl-oleanolic acid. Furthermore, CaCYP716E41 can also catalyze maslinic acid to produce 6β-hydroxy maslinic acid (Miettinen et al., 2017). To investigate whether NaCYP716E107 and SlCYP716E26 can accept other triterpene scaffolds, we supplemented leaves expressing NaCYP716E107 or SlCYP716E26 with substrates such as erythrodiol (N2), oleanolic acid (N4), or maslinic acid (N7). No new peaks were detected when erythrodiol (N2) and maslinic acid (N7) were added to the leaves (**Fig. 5C**). However, upon the addition of oleanolic acid (N4), a new peak, N9, was detected in leaves expressing either NaCYP716E107 or SlCYP716E26, albeit in a low yield (**Fig. 5C, Fig. S17B**). We hypothesized that the efficiency of substrate uptake into plant cells might be too low to provide sufficient substrate for the enzymes. Therefore, we supplemented extracts of NaOSC2/NaCYP716A419 expressing leaves to produce oleanolic acid and extracts of NaOSC2/NaCYP716A419/CYP716C87 expressing leaves to produce maslinic acid *in vivo* (**Fig. 5D**). Co-expression of NaOSC2/NaCYP716A419 with NaCYP716E107 or SlCYP716E26 resulted in N9 in greater abundance and another new peak, N10 (**Fig. 5D, Fig. S5**).

**Fig. 5.**
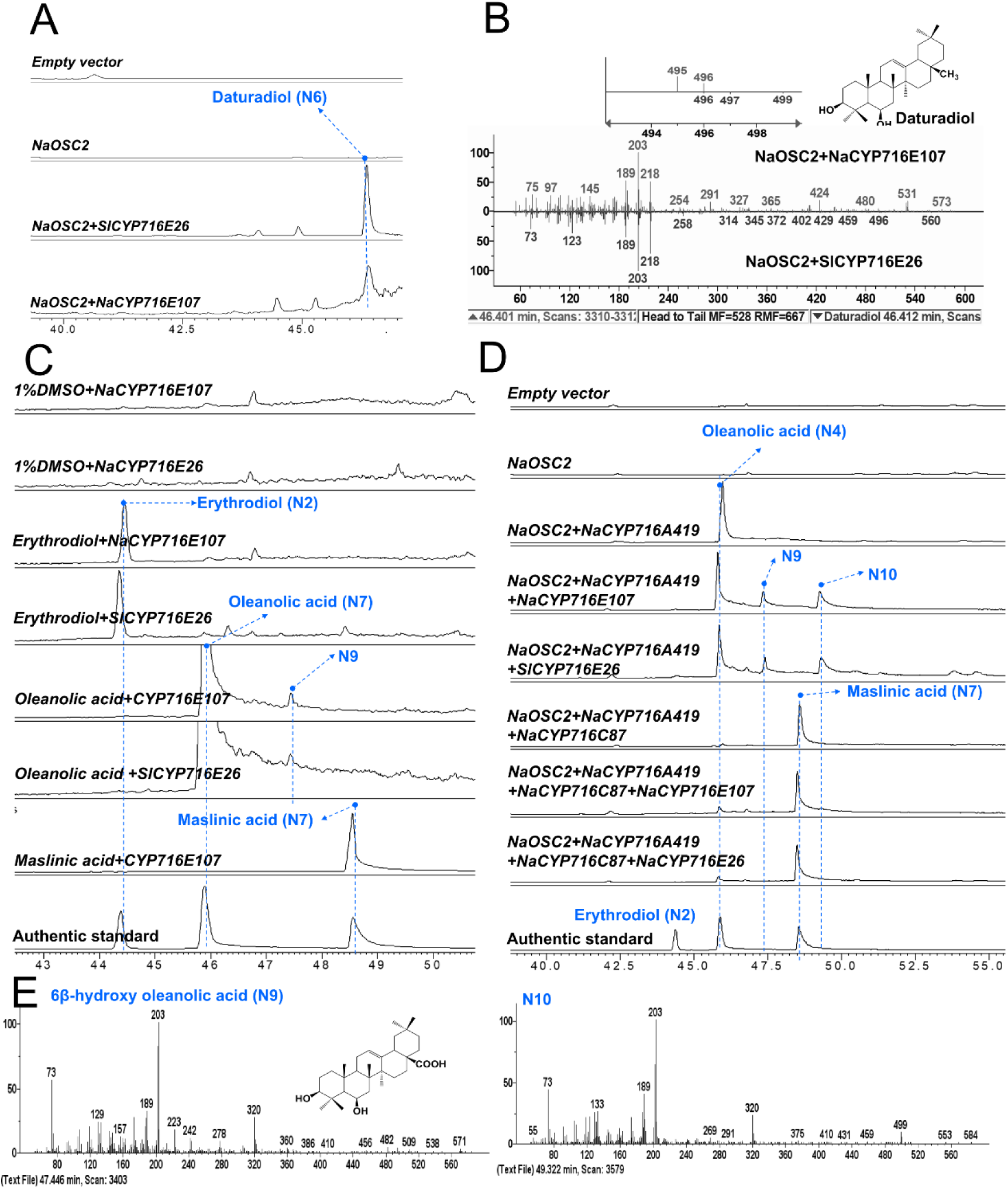
CYP716E107 is a C6β oxidase. **(A)** GC-SIM MS (m/z: 189, 203, 218) profile of extracts of *N. benthamiana* leaves co-expressing NaOSC2/CYP716E107 (SlCYP716E26). SlCYP716E26 is a P450 monooxygenase cloned from *Solanum lycopersicum*, responsible for the C6β oxidation of β-amyrin (Yasumoto et al., 2017). **(B)** GC-MS spectrum of daturadiol. **(C)** GC-SIM MS (m/z: 189, 203, 320) profiles of extracts of *N. benthamiana* leaves feed with different substrates. **(D)** GC-SIM MS (m/z: 189, 203, 320) profiles of extracts of *N. benthamiana* leaves co-expressing NaCYP716E107 or SlCYP716E26 with NaOSC2/NaCYP716A419 or NaOSC2/NaCYP716A419/NaCYP716C87. **(E)** EI-MS spectra of trimethylsilylated 6β-hydroxy-oleanolic acid (N9), and N10. Each treatment was conducted with at least three replications.

Based on the comparison of the EI-MS spectra of N10 with the fragmentation pathway of oleanolic acid (**Fig. 5E and Fig. S10**), as well as the similarity of N9 and N10 to the products of CaCYP716E41 reported in (Miettinen et al., 2017), we speculate that N9 is 6β-hydroxylated oleanolic acid, and N10 is incompletely derivitized 6β-hydroxylated oleanolic acid. However, even the expression of NaCYP716E107 or SlCYP716E26 with the co-expression of NaOSC2/NaCYP716A419/CYP716C87, did not produce detectable quantitites of 6β-hydroxyl-maslinic acid (**Fig. 5D, Fig. S5**). We also attempted co-expression of NaCYP716E107 with AtLUP1 or AtLUP1/NaCYP716A419 but did not observe any new peaks (**Fig. 1, Fig. S3 and Fig. S6**).

### Expression patterns of Na*cyp716a419*, Na*cyp716e107*, and Na*cyp716c87*

To investigate the potential functions of these CYP716 enzymes in *N. attenuata*, we conducted real-time quantitative PCR (RT-qPCR) analysis to examine their expression and induction patterns. Consistent with the results obtained from the microarray analysis in **Fig. 1**, the relative transcript levels of *Nacyp716a419*, *Nacyp716e107*, and *Nacyp716c87* were found to be higher in flowers, followed by roots, with the lowest expression observed in leaves. Notably, the expression level of the *Nacyp716c87* gene in roots was comparable to that in flowers (**Fig.6A**). Additionally, we investigated the responses of three *Nacyp716a419*, *Nacyp716e107*, and *Nacyp716c87* to various phytohormones and abiotic stresses, including abscisic acid (ABA), salicylic acid (SA), gibberellin A3 (GA_3_), and methyl jasmonate (MeJA), as well as PEG6000 (mimicking drought stress), NaCl (salt stress), and Na_2_CO_3_ (mimicking alkaline stress) (**Fig. 6B-E**). After 4h treatments with ABA, SA, GA_3_, and MeJA, the relative transcript levels of *Nacyp716a419* in the roots increased by 1.7, 6.5, 15.3, and 4.9-fold, respectively, compared to the control group (**Fig. 6B**). In contrast, *Nacyp716e107* exhibited a weaker response to these four phytohormones. ABA, GA_3_, and MeJA induced *Nacyp716e107* most strongly 7h after treatment, resulting in a 4-fold increase in relative transcript levels for ABA, a 3.5-fold increase for GA_3_, and a 3-fold increase for MeJA treatments, compared to the control. No significant induction of *Nacyp716c87* transcripts was observed within the first 9h of MeJA treatment. However, 4h after treatments with ABA, SA, and GA_3_, the relative transcript levels of *Nacyp716c87* increased by 6.1, 117, and 41.7-fold, respectively, compared to the control group (**Fig. 6B**). In response to PEG6000 treatments, simulating drought stress, *Nacyp716e107*, and *Nacyp716c87* transcript levels increased 3- and 5-fold, respectively, compared to the control group (0% PEG) (**Fig. 6C**). In contrast, *Nacyp716a419* was largely unresponsive. In response to NaCl treatments, relative transcript levels of *Nacyp716a419*, *Nacyp716e107*, and *Nacyp716c87* all showed reductions when compared to the control group (0 mM NaCl) (**Fig. 6D**). Specifically, treatments with 50 and 100 mM NaCl resulted in 19 and 37% decreases in *Nacyp716a419*, 25 and 36% decreases in *Nacyp716e107*, and 94 and 97% decreases in *Nacyp716c87* relative transcript levels. Under alkaline stress (Na_2_CO_3_ treatment), only transcripts of *Nacyp716a419* and *Nacyp716e107*increased significantly. Specifically, at 100 and 200 mM Na_2_CO_3_ treatments, *Nacyp716a419* increased 10- and 17-fold, respectively, and *Nacyp716e107* increased 41- and 56-fold, respectively (**Fig. 6E**)., *Nacyp716a419*, *Nacyp716e107*, and *Nacyp716c87* appear to be responsive to a variety of environmental stressors in *N. attenuata*. The response of *Nacyp716a419*, *Nacyp716e107*, and *Nacyp716c87* to hormones and abiotic stresses exhibits notable differences, suggesting they may possess distinct biological functions. *Nacyp716a419*, *Nacyp716e107*, and *Nacyp716c87* all show a response to GA_3_ and ABA or MeJA treatments, indicating their potential involvement in growth-related or defense-related processes.

**Fig. 6.**
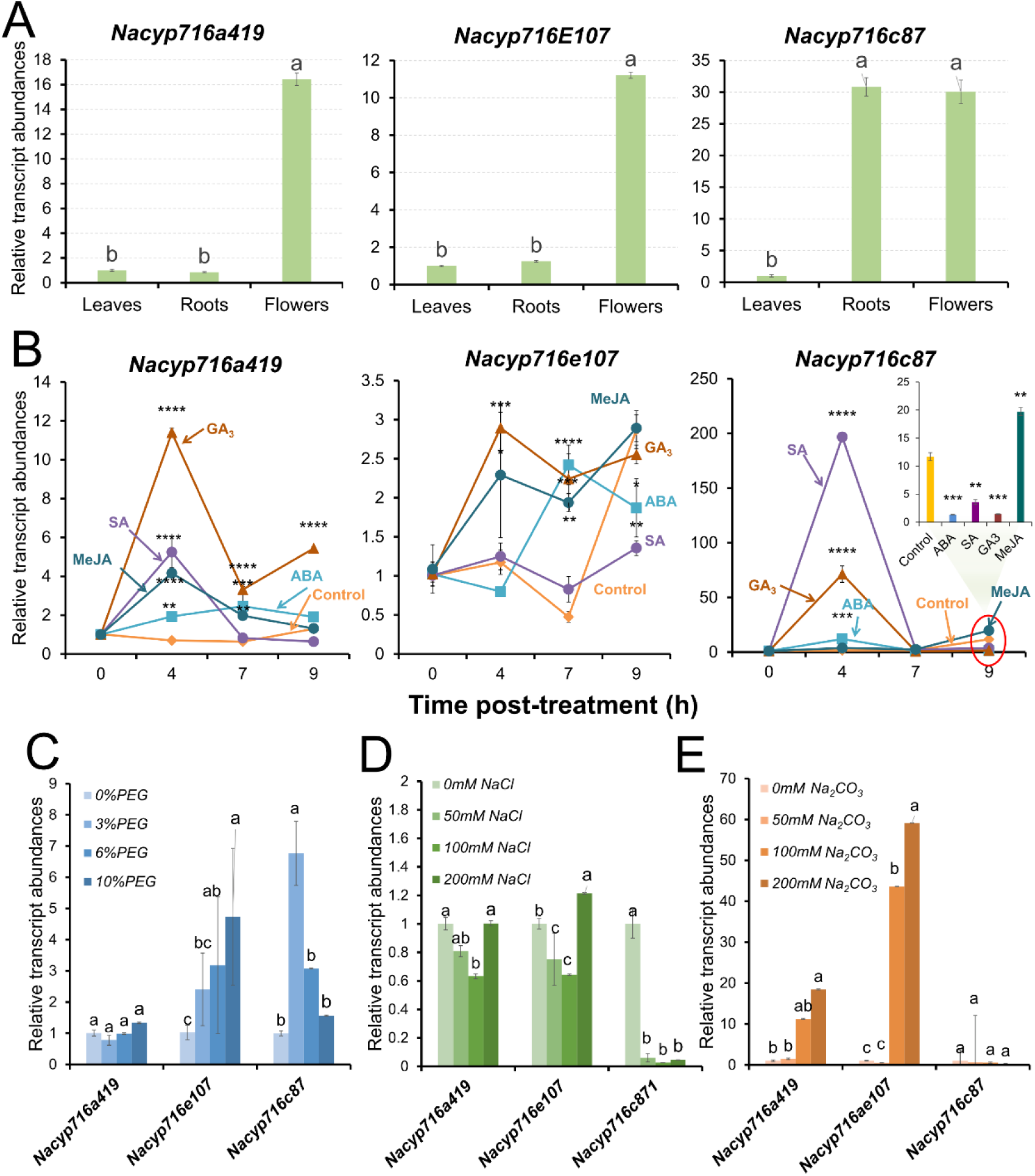
Responses of *CYP716A419*, *CYP716E107*, and *CYP716C87* transcripts to hormone and abiotic stress treatments. **(A)** Relative transcript levels of *Nacyp716a419*, *Nacyp716e107*, and *Nacyp716c87* in different tissues. () Relative transcript levels of *CYP716A419*, *CYP716E107*, and *CYP716C87* after phytohormone treatment. Inset provides statistical details of the 9h harvest for *CYP716C87.* **(B)** Relative transcript levels of *CYP716A419*, *CYP716E107*, and *CYP716C87* after PEG6000 treatments. **(C)** Relative transcription levels of *CYP716A419*, *CYP716E107*, and *CYP716C87* after NaCl treatment. **(D)** Relative transcription levels of *CYP716A419*, *CYP716E107*, and *CYP716C87* after Na_2_CO_3_ treatment. Results of ANOVAs with Tukey’s test are shown in panels A and C (n=3, mean ± SE; different lowercase letters indicate statistically significant differences between two groups of samples at the *p* < 0.05 level). Results of Student’s t-tests are shown in panel B,(*, *p* < 0.05; **, *p* < 0.01; ***, *p* < 0.001; ****, *p* < 0.0001).

### Silencing *Nacyp716a419* attenuates *N. attenuata* growth and reproductive performance

To further elucidate the biological functions of *Nacyp716a419*, *Nacyp716c87*, and *Nacyp716e107* in *N. attenuata*, we employed a virus-induced gene silencing (VIGS) technique using Tobacco Rattle Virus (TRV) (Saedler and Baldwin, 2004). Each of the CYP450 genes was individually silenced in separate replicate plants. Additionally, plants inoculated with TRV vectors harboring empty vector (EV) constructs were included as negative controls and *NaOSC1-* and *NaOSC2-*harboring VIGS vectors served as positive controls (*NaOSC1/2_VIGS*). The visibly apparent bleaching phenotype of tomato phytoene desaturase (PDS) -silenced plants (*PDS_VIGS*) were used to monitor the onset and spread of gene silencing.

Given that most plant-specialized metabolites function in defense or stress responses, phenotyping of plant growth under intense competition and nutrient limitations provides a useful means of amplifying the growth-related costs associated with metabolite production (Baldwin and Hamilton, 2000; Yang et al., 2023). Hence, a companion-plant approach was used, in which two plants matched in initial size were grown in a single pot, with one plant silenced in the expression of different triterpene biosynthesis genes (*Naosc1/2, Nacyp716a419, Nacyp716e107,* or *Nacyp716c87*) and the other was a control EV-inoculated plant. This setup facilitates the quantification of subtle differences in plant growth and reproductive performance resulting from silencing these triterpene biosynthesis genes. Each combination was performed with 30 replicates. Within 14 days after *Agrobacterium* -infiltration, plants that did not exhibit viral infection symptoms were discarded. At 25 days post *Agrobacterium* infiltration, newly grown leaves of *PDS_VIGS* plants became completely bleached, indicating the successful induction of virus-mediated gene silencing (**Fig. 7A**). qPCR analysis of VIGS plant leaves revealed that, compared to EV plants, *NaOSC1/2-VIGS* plants had relative transcript levels reduced by 56 and 44% for *Naosc1* and *Naosc2*, respectively. *Nacyp716a419* silenced plants (*CYP716A419_VIGS*) exhibited a 45% reduction in *Nacyp716a419* transcript levels, while *Nacyp716e107*silenced plants (*CYP716E107_VIGS*) plants showed a 74% reduction in *Nacyp716e107* transcript levels. Na*cyp716e107*silenced plants (*CYP716C87_VIGS*) plants displayed a 48% reduction in *Nacyp716e107*transcript levels (**Fig. S18**). After 34 days of *Agrobacterium* inoculation, we observed significant growth inhibitions in *CYP716A419_VIGS* plants (**Fig. 7A**). This inhibition was evident in reduced stalk height and rosette diameter **Fig. 7B** compared to the neighboring EV plants (**Fig. 7A**). As the plants entered the flowering stage, these growth differences became more pronounced. In *CYP716A419_VIGS* plants, the leaves became slender, and flower buds were aborted (**Fig. 7A**). Statistical analysis revealed that rosette diameters of *CYP716A419_VIGS* plants were 24% smaller than those of both neighboring EV plants and the EV-EV plants (**Fig. 7B**). Furthermore, *CYP716A419_VIGS* plants were 33% thinner in stalk diameter, 29% smaller in stalk height, and matured 59% fewer seed capsules compared to their neighboring EV plants. These reductions were 31%, 29%, and 51%, respectively, of those of the EV-EV control group (**Fig. 7B**). While significant reductions in rosette diameters and stalk heights were also observed in plants silenced in *Nacyp716e107*, and silencing *Naosc1/2* resulted in a notable reduction in rosette diameters compared to EV plants.These plants did not show significant differences in capsule numbers compared to the EV controls (**Fig. 7B**).Due to the poor reproductive performance of CYP716A419_VIGS plants in terms of capsule numbers, we evaluated the pollen viability of these plants. Metabolic staining of pollen revealed lower pollen viability in CYP716A419_VIGS plants compared to both EV plants and NaOSC1/2_VIGS plants (**Fig. S19**).

**Fig. 7.**
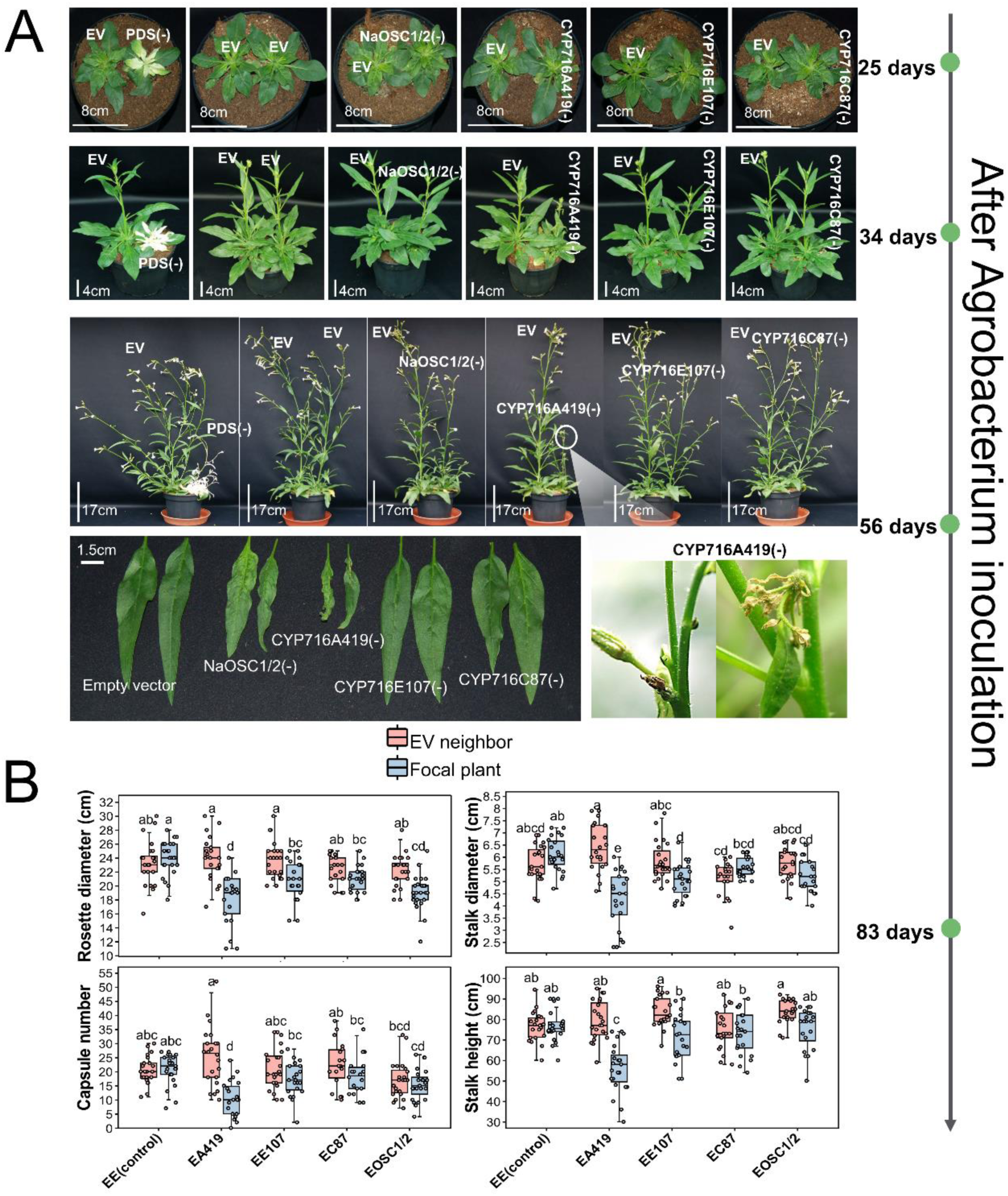
Silencing key triterpenoid biosynthesis genes attenuate the growth and fitness of *N. attenuata*. **(A)** The growth phenotypes of VIGS plants after *Agrobacterium* inoculation. Only *CYP716A419_VIGS* plants aborted flowers. **(B)** Fitness of VIGS plants (*NaOSC1/2_VIGS, CYP716A419_VIGS, CYP716E107_VIGS* or *CYP716C87_VIGS*). Focal plants refer to the plants in which *Naosc1/2*, *Nacyp716a419*, *Nacyp716e107*, or *Nacyp716c87* were silenced. Neighbor plants refer to initially size-matched plants inoculated with *Agrobacterium* carrying an empty vector (EV) construct. The control group, labeled EE(control), consists of two EV plants. EA419 indicates that the neighbor plants are EV plants, and the focal plants are *CYP716A419_VIGS* plants. Similar labeling schemes for the other combinations: EE107: EV and *CYP716E107-VIGS* plant pairs; EC87: EV and *CYP716C87_VIGS* plant pairs; EOSC1/2: EV and *NaOSC1/2_VIGS* plants pairs. Results of two-tailed ANOVAs with Tukey’s tests are shown (n= 21∼24, mean ± SE, different lowercase letters indicate statistically significant differences between two groups of samples at the *p* < 0.05 level). The central line within the box represents the median of the data. The upper and lower boundaries of the box denote the upper and lower quartiles of the data. The lines above and below the box, known as whiskers, signify the variability of the data (error bar). The whiskers’ length is set at 1.5 times the interquartile range. The points represent specific observations in the data set. The values beyond 1.5 times the interquartile range are considered as outliers.

### Triterpenoids in *N. attenuata*

Given the abundant transcript levels of enzymes NaCYP716A41, NaCYP716E107, and NaCYP716C87 in floral organs, our analysis focused on triterpenoids in floral parts of VIGS plants. In the floral organs of VIGS plants, β-amyrin and the products of NaCYP716A419 (erythrodiol and oleanolic aldehyde) were detected (**Fig. 8**).

**Fig. 8.**
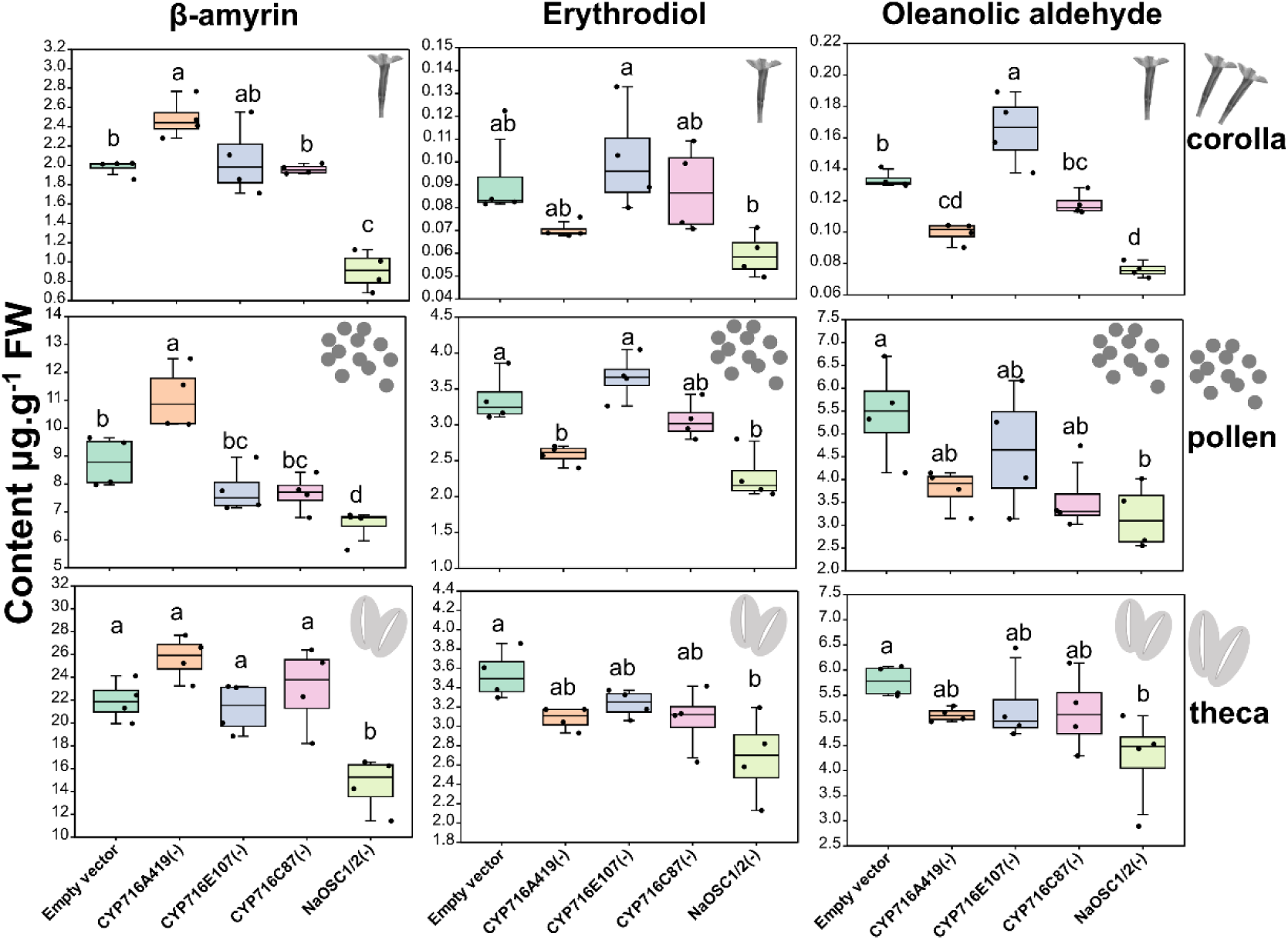
Triterpene contents in CYP450-silenced plants. **(A)** Triterpene contents in flower parts.. Samples from 4 individual plants were mixed to form a single replicate; a total of 4-6 replicates for each tissue type were used. ANOVAs with Tukey’s tests are shown (n=4∼6, mean ± SE, different lowercase letters indicate statistically significant differences between two groups of samples at the *p* < 0.05 level). The central line within the box represents the median of the data. The upper and lower boundaries of the box denote the upper and lower quartiles of the data. The lines above and below the box, known as whiskers, signify the variability of the data (error bar). The whiskers’ length is set at 1.5 times the interquartile range. The points represent specific observations in the data set. The values beyond 1.5 times the interquartile range are considered as outliers.

Silencing *Nacyp716a419* led to a 26% increase in β-amyrin content in the corolla, 26% in the pollen, and 17% in the thecal coverings (theca) of the anther heads which enclose the pollen grains until dehiscence. These increases were associated with decreases in erythrodiol, by 24, 23, and 13%, and oleanolic aldehyde, by 26, 26, and 12%, in these respective floral tissues. In the case of *Nacyp716e107* silencing, only oleanolic aldehyde increased by 25% in the corolla, while the other components remained unchanged. In *Naosc1* and *Naosc2* co-silenced plants, there were significant reductions in triterpenoid contents when compared to the EV control. Specifically, β-amyrin decreased by 54% in the corolla, 26% in the pollen, and 33% in the theca.

Erythrodiol showed a 36% reduction in the corolla, a 32% reduction in the pollen, and a 24% reduction in the theca. Meanwhile, oleanolic aldehyde decreased by 43% in the corolla, 42% in the pollen, and 27% in the theca (**Fig. 8**). Notably, these compounds did not exhibit significant changes in plants with *Nacyp716c87* silenced (**Fig. 8**).

We did not detect maslinic acid, 2α-hydroxyβ-amyrin (products of NaCYP716C87), or 6β-hydroxy oleanolic acid (products of NaCYP716E107) in the flowers of *N.attenuata*. To explore whether the enzyme products accumulate in *N. attenuata*, and considering that ABA, GA_3_, SA, and MeJA can induce the expression of the *Nacyp716c87*, *Nacyp716a419*, and *Nacyp716E107* genes in the root, we attempted to detect the triterpenoids in plant roots 72h after induction with ABA, GA_3_, SA, and MeJA. No products of NaCYP716C87 and NaCYP716E107 were observed; instead, peaks suspected to be derivatives of lupanediol were detected in these hormone-induced plants (**Fig. S20**). However, it is noteworthy that oleanolic acid (N4), maslinic acid (N7), and 6β-hydroxy oleanolic acid (N9) were detected in the roots colonized for 5 weeks by arbuscular mycorrhizal fungi (AMF) (**Fig. S21**). These results suggest that NaCYP716A419, NaCYP716C87, and NaCYP716E107 can produce several of their respective products in *N. attenuata*, but their production depends on specific developmental stages and specific inducing factors.

## Discussion

In this study, we characterized the functions of several P450 enzymes from *N. attenuata* in *N. benthamiana* (**Fig. 1**). The work revealed that NaCYP716A419, NaCYP716C87, and NaCYP716E107 modify pentacyclic triterpene skeletons, specifically at the C28, C2α, and C6β positions of β-amyrin, lupeol, or lupanediol, respectively.

The CYP716A subfamily has been extensively studied in other plants, and is known to catalyze a sequential three-step oxidation at the C28 position of β-amyrin, lupeol, and α-amyrin, forming a hydroxyl group, aldehyde, and carboxyl group, respectively (Carelli et al., 2011; Fukushima et al., 2011; Miettinen et al., 2017; Misra et al., 2017; Yasumoto et al., 2017; Alagna et al., 2023). Certain members of the CYP716A subfamily, such as BpCYP716A180 from *Betula platyphylla*, have been suggested to catalyze C28 oxidations on lupanediol (Zhou et al., 2016).

NaCYP716A419, when co-expressed with NaOSC2 in *N. benthamiana*, yielded erythrodiol (N2), oleanolic acid (N4), and oleanolic aldehyde (N5). Additionally, when co-expressed with AtLUP1 in *N. benthamiana*, the enzyme resulted in the production of betulin (A3), betulinic acid (A4), 28-hydroxy lupanediol (A6), and 28- carboxy lupanediol (A8), indicating that NaCYP716A419 also functions as a C-28 oxidase.

In contrast to the well-characterized CYP716A subfamily enzymes, the enzymatic function of the CYP716C subfamily remains largely unknown. To date, only CYP716C11 from *Centella asiatica* (Miettinen et al., 2017), CYP716C53 from *Avicennia marina* (Nakamura et al., 2018), CYP716C67 from *Olea europaea* (Alagna et al., 2023), and CYP716C49 from *Crataegus pinnatifida* (Dai et al., 2019), as well as CYP716C55 from *Lagerstroemia speciosa* (Sandeep et al., 2019), have been characterized. These enzymes catalyze the hydroxylation of the C2α position of oleanolic acid, ursolic acid, or betulinic acid, thereby forming maslinic acid, corosolic acid, and alphitolic acid (Miettinen et al., 2017; Nakamura et al., 2018; Dai et al., 2019; Alagna et al., 2023). However, there have been limited studies indicating the activity of CYP716C subfamily members on other triterpene alcohol substrates.

NaCYP716C87 catalyzes the hydroxylation of the C2α position of oleanolic acid and betulinic acid, indicating that NaCYP716C87 is also a C2α hydroxylase. When NaCYP716C87 is co-expressed with NaOSC2 or AtLUP1 in *N. benthamiana* (**Fig. 1**), it yields 2α-hydroxy β-amyrin (N3), 2α-hydroxy lupeol (A2), and A7 (2α-hydroxy lupanediol). Feeding erythrodiol to the leaves expressing NaCYP716C87 results in the production of 2α-hydroxy erythrodiol. These results suggested that some CYP716C family members can also catalyze C2α hydroxylation of triterpene alcohols.

Many members of the CYP716E subfamily have not yet been functionally characterized. To date, only two members, SlCYP716E26 and CaCYP716E41, have been reported to participate in the biosynthesis of triterpenoids (Miettinen et al., 2017; Yasumoto et al., 2017). In this study, NaCYP716E107 was found to act on the β- amyrin and oleanolic acid skeletons, but not on lupeol or lupanediol (**Fig. 5**). Previous reports showed that SlCYP716E26 catalyzes hydroxylation at the C6β position of β- amyrin, forming daturadiol (N6), but its activity towards other substrates was not investigated (Yasumoto et al., 2017). Conversely, CaCYP716E41 was reported to not catalyze β-amyrin but to catalyze hydroxylation at the C6β position of oleanolic acid and maslinic acid (Miettinen et al., 2017). In our study, we cloned SlCYP716E26 from tomato and compared it with the products of NaCYP716E107, demonstrating that NaCYP716E107 also functions as a C6β hydroxylase, catalyzing β-amyrin to form daturadiol (N6). Additionally, both NaCYP716E107 and SlCYP716E26 can catalyze oleanolic acid, consistent with CaCYP716E41’s function, producing two products (putative 6β-hydroxy-oleanolic acid and its incompletely derivitized product), but they cannot catalyze maslinic acid substrates, which CaCYP716E41 can. These differences may be related to substantial variations in their protein sequences at positions 251 to 260 (**Fig. S16**).

In this study, other enzymes (CYP716A420, CYP716E108, CYP716D93, CYP716D94, CYP716H6, CYP88B4, and CYP88C16) did not show activity towards β-amyrin or lupeol (**Fig. 1**). We attempted co-expression of these inactive enzymes with NaCYP716A419, NaCYP716C87, or NaCYP716E107 in *N. benthamiana*, but still no products were detected (**Fig. S22 and Fig. S23**). Given that some CYP450 enzymes only accept substrates that have been modified with specific substituents, we cannot rule out the possibility that these enzymes may be involved in triterpene biosynthesis. For instance, MtCYP72A61v2 and MtCYP72A68v2 exhibit C-22β and C-23 oxidation activity, respectively, towards 24-hydroxy-β-amyrin and ursolic acid, but they do not catalyze the oxidation of β-amyrin (Fukushima et al., 2013).

GuCYP72A154 from *G. uralensis* is a C30 oxidase capable of catalyzing the three-step oxidation of 11-oxo-β-amyrin to glycyrrhizin instead of directly oxidizing the β- amyrin scaffold (Seki et al., 2011). Furthermore, some members of the CYP716 family can catalyze the oxidation of tetracyclic triterpenes (Ghosh, 2017). For example, ginseng’s GuCYP716A47 (renamed CYP716U1) catalyzes the conversion of dammarenediol-II into protopanaxadiol (Han et al., 2011), and GuCYP716A53v2 (renamed CYP716S1v2) catalyzes the transformation of protopanaxadiol into protopanaxatriol (Han et al., 2012). CsCYP88L2 and CsCYP81Q58 from *Cucumis sativus* catalyze two consecutive oxidation reactions in the cucurbitacin biosynthesis pathway (Shang et al., 2014). Our previous research has also confirmed the presence of dammarenediol, taraxasterol, and several other unidentified triterpene scaffolds in *N. attenuata* (Yang et al., 2023). Hence, we cannot exclude the possibility that CYP716A420, CYP716E108, CYP716D93, CYP716D94, CYP716H6, CYP88B4, and CYP88C16 may exhibit catalytic activity towards these triterpenes.

*Nacyp716a419*, *Nacyp716c87*, and *Nacyp716e107* exhibit the highest expression in flowers. However, in the flowers of VIGS plants in which these enzymes were individually knocked down, only in NaCYP716A419 VIGS plants, could erythrodiol and oleanolic aldehyde be detected in modest decreases: we did not observe the enzyme products of NaCYP716C87 and NaCYP716E107. Given the transcript elicitations of these enzymes by GA_3_, ABA, and MeJA treatments in roots (**Fig. 6B**), we also attempted to detect the NaCYP716C87 and NaCYP716E107 triterpenoid products after hormone inductions, but without success (**Fig. S20**). Metabolomic analysis suggests the possible existence of glycosylated compounds derived from lupeol or β-amyrin in *N. attenuata* (Yang et al., 2023). The enzyme products identified in this study are intermediates; hence, the enzyme products of NaCYP716C87 and NaCYP716E107 may have been metabolized into downstream compounds, thwarting the detection of the intermediates. Additionally, detectable levels of oleanolic acid (N4), maslinic acid (N7), 6β-hydroxy-oleanolic acid (N9), and N10 were found in roots after AMF colonization. This indicates that these compounds may be detected in *N. attenuata* only under specific growth conditions and this provides an alternative explanation for lack of product detection for NaCYP716C87 and NaCYP716E107..

We also observed that CYP716A419-VIGS plants exhibited a clear reductions in capsule numbers, as well as decreased stem height, diameter, and rosette diameter, along with attenuated pollen viability. However, silencing the initial enzyme of the triterpenoid biosynthetic pathway, NaOSC1/2-VIGS plants did not result in similarly severe effects. In a previous study in *N. attenuata*, we observed a similar phenomenon with a different biosynthetic pathway. Silencing the specific geranylgeranyl pyrophosphate (GGPPS) that controls diterpene biosynthetic flux into 17- hydroxygeranyllinalool diterpene glycoside biosynthesis, did not result in developmental abnormalities. However, silencing genes involved in later steps in this pathway, such as *UGT74P3*, *UGT74P5*, *CYP736A304*, or *CYP736A305*, led to the accumulation of 17-hydroxygeranyllinalool or geranyllinalool, resulting in severe developmental defects (Heiling et al., 2021; Li et al., 2021). In many plants, intermediates of triterpene biosynthesis accumulate during normal growth and development, and manipulating their biosynthesis can lead to morphological and physiological effects (Guhling et al., 2006; Ohyama et al., 2007). For instance, *Arabidopsis thaliana* plants that overexpress thalianol synthase (THAS) are dwarfed but produce longer roots (Field and Osbourn, 2008). Mutants of the marneral synthase (MRN1) gene in *Arabidopsis* display delayed embryogenesis, late flowering, rotund leaves, and abnormal seed morphology (Field et al., 2011). In oats, an increase in β- amyrin in the roots leads to shorter roots with a super-hairy phenotype (Kemen et al., 2014). In addition, disrupting the oxidation, glycosylation, or acylation steps in triterpene synthesis can also lead to developmental phenotypes. For example, the loss of function mutants in genes MtCYP716A12 and UGT73F3 in *Medicago truncatula* truncates the production of hemolytic saponins and results in a dwarfing phenotype (Naoumkina et al., 2010; Carelli et al., 2011). Some triterpenoid saponins can regulate growth-related hormones, such as chromosaponin I (CSI), a γ-pyronyl-triterpenoid saponin isolated from peas (*Pisum sativum*) and other leguminous plants (Kudou et al., 1992; Tsurumi et al., 1992), specifically interacts with the auxin influx carrier AUX1 thereby altering responses of *Arabidopsis* roots to auxin and ethylene (Rahman et al., 2001). Hence, we hypothesize that the observed phenotypic alterations could be attributed to either the toxic effects arising from the accumulation of intermediate products due to the silencing of Nacyp716a419 or the absence of downstream growth-related triterpenoid compounds.

In summary, heterologous expression in *N. benthamiana* revealed that NaCYP716A419, NaCYP716E107, and NaCYP716C87 catalyze the oxidation of β- amyrin, lupeol, lupanediol, or their downstream compound skeletons at C28, C6β, and C2α positions. We attempted to validate these enzyme functions in *N. attenuata* by silencing *Nacyp716Aa19*, *Nacyp716e107*, and *Nacyp716c87*. We attempted to detect these enzyme products in hormone- or stress-induced *N. attenuata* roots. We found maslinic acid (N7), 6β-hydroxy-oleanolic acid (N9), and N10 in AMF-harboring roots. While their concentrations were low, these products clearly accumulate in *N. attenuata*. Additionally, our study indicated that CYP716A419-VIGS plants exhibited significant growth inhibition and reduced capsule numbers under competitive conditions. However, due to the transient and unstable silencing of VIGS, we can only conclude that NaCYP716A419 likely has a notable impact on the growth of *N. attenuata*. The specific functions and regulatory mechanisms of how NaCYP716A419 influences growth and development of *N. attenuata* will require additional work with stable gene knockouts.

## MATERIALS AND METHODS

### Chemicals

Methyl jasmonate (product ID: W341002), abscisic acid (product ID: A4906), salicylic acid (product ID: 84210), α-amyrin (product ID: 53017), β-amyrin (product ID: 09236), erythrodiol (product ID: 09258), lupeol (product ID: 18692), betulinic acid (product ID: 91466), oleanolic acid (product ID: 42515), N-Methyl-N-trimethylsilyl-trifluoroacetamide (MSTFA) (product ID: 69479), and N, N-Dimethylformamide (DMF) (product ID: 09258), were purchased from Sigma-Aldrich (St Louis, MO, USA), and gibberellin A_3_ (product ID: Art. No. 7464.2) was purchased from Carl Roth (Karlsruhe, Germany).

### Cultivation and elicitation of *N. attenuata* plants

*N. attenuata* Torr. Ex Watts seeds from the 31^st^-generation inbred line were utilized as the wild-type (WT) genotype in all experiments. Seed germination and plant growth followed the protocols outlined in previous reports (Krügel et al., 2002), with a day/night cycle of 16h (26-28℃) and 8h (22-24℃) in a glasshouse at the Max Planck Institute for Chemical Ecology, Jena, Germany.

For phytohormone and abiotic stress inductions, 10-day-old seedlings were transferred to a substrate composed of soil balls and sand in a 1:3 ratio for 3 weeks. 20 μL of lanolin containing 100 μM methyl jasmonate, abscisic acid, salicylic acid, or gibberellin A_3_ were applied to the basal ends of seedling stems. An equivalent amount of pure lanolin served as a negative control. Roots were collected at 0, 4, 7, and 9 h after treatments for transcript analysis. Roots were collected after 72 h treatments for triterpenoids analysis. For abiotic stress treatments (salt, alkali, and drought stress), 50 mL of different concentrations of NaCl (0, 50, 100, 200 mM), Na_2_CO_3_ (0, 50, 100, 200 mM), and PEG6000 (0, 3%, 6%, 10%) were added to the sand to create salt, alkali, and drought stress conditions, respectively. After 2 days of stress-treatments, root samples were harvested for transcript analysis.

### Phylogenetic analysis and sequence alignment

The protein sequences of cytochrome P450 enzymes were obtained from the NCBI GenBank (https://www.ncbi.nlm.nih.gov/genbank/) and the *Nicotiana attenuata* data hub (Brockmöller et al., 2017). Details regarding cytochrome P450 enzymes related to triterpene biosynthesis are provided in Supplementary Table S1. Sequence alignment was conducted using ClustalW, and a phylogenetic tree was constructed through MEGA11 (Tamura et al., 2021) employing the Neighbor-Joining method with 1000 bootstrap replicates. The tree’s aesthetic enhancement was accomplished using tvBOT (Xie et al., 2023).

### Heterologous expression of candidate enzymes in *Nicotiana benthamiana*

Full-length cDNAs of selected cytochrome P450 and NaOSC2 enzymes from *N. attenuata, Atlup1* from *A. thaliana,* and CYP716E26 from *S. lycopersicum* were cloned into a 3Ω1 expression vector (Cárdenas et al., 2019; Hong et al., 2022) using the ClonExpress® II One Step Cloning Kit (Vazyme) with the primers listed in **Supplementary Table S5** and the vectors were transformed into *Agrobacterium tumefaciens* strain *GV3101*. A heterologous expression followed established protocols (Yang et al., 2023). *Agrobacterium* strains carrying the gene constructs were cultured in 10 mL of LB medium supplemented with antibiotics (100 μg·mL^-1^ rifampicin and 250 μg·mL^-1^ spectinomycin) at 28 °C for 24 h. After centrifugation, the supernatant was discarded, and the cell pellet was resuspended in 5 mL of infiltration buffer (50 mM MES, 2 mM Na_3_PO_4_, 10 mM MgCl_2_, and 100 μM acetosyringone). The cell suspension was then diluted to an OD600 of 0.6 for single enzymes and 0.4 for enzyme combinations. Infiltration was performed on the abaxial surface of 4-5-week-old *N. benthamiana* leaves using a needle-free syringe. Triterpenoid analysis was conducted five days after *Agrobacterium* infiltration, and leaves were collected from three individual plants. For substrate feeding, 1 mL of 100 µM substrates (dissolved in 1% DMSO) was infiltrated into the abaxial surface of leaves three days after *Agrobacterium* inoculation. Leaf samples were collected for triterpenoid analysis three days after substrate injection.

### Virus-induced gene silencing (VIGS)

VIGS experiments were performed according to a published protocol optimized for VIGS in *N. attenuata* (Saedler & Baldwin, 2004) with the *pBINTRA* and *pTV00* VIGS expression system. Briefly, 260 to 300bp mRNA (CDS or UTR) sequences of enzymes were amplified by PCR using Q5® High-Fidelity DNA Polymerase (New England Biolabs) with primers listed in **Supplementary Table S6**and cut by BamHI and SalI (New England Biolabs). The DNA fragments were cloned into a *pTV00* vector using T4 DNA ligase (Promega) and transformed into *Agrobacterium tumefaciens strain GV3101*.

Two size-matched 20-day-old seedlings were transplanted into individual 2 L pots and placed in a controlled environment chamber with a 16/8-h day/night cycle (26-28°C during the day, 22-24°C at night) and 40% relative humidity. Four days after transplantation, one of the seedlings from each pair was inoculated with *Agrobacterium* harboring the pBINTRA vector along with a VIGS vector containing gene fragments related to triterpene biosynthesis (*pTV-OSC1/2*, *pTV-CYP716A419*, *pTV-CYP716E107*, or *pTV-CYP716C87*). Thirty plants were inoculated with each vector. The neighboring plants of these individuals were inoculated with *Agrobacterium* containing empty vectors (EV) and served as negative controls within each group. Pots with two EV plants were used as negative controls. Plants inoculated with *Agrobacterium* carrying the phytoene desaturase (PDS) gene served as positive controls and the onset of leaf-bleaching was used to time leaf harvest. Fourteen days after *Agrobacterium* inoculation, plants without apparent viral phenotypes were removed from the study. Each experimental group retained 21 to 24 replicate plants.

### Transcript abundance analysis

The total RNA was extracted from the plant tissues of *N. attenuata* by utilizing the plant RNA purification kit (Macherey-Nagel) following the manufacturer’s instructions. Complementary DNA (cDNA) was generated from total RNA using PrimeScript RT Master Mix (Takara Bio Inc., Japan). RT-qPCR was performed on a Stratagene Mx3005P qPCR machine using a Takyon™ No ROX SYBR 2X MasterMix Blue dTTP (Eurogentec, Seraing, Belgium). The housekeeping gene IF-5α from *N. attenuata* was used as an internal reference. The primers used for RT-qPCR are listed in **Supplementary Table S7**.

### GCMS analysis of triterpenes

Triterpene extraction followed established procedures outlined in a prior publication (Field and Osbourn, 2008) with minor modifications. For triterpene extraction from roots, leaves, and corollas of *N. attenuata*, approximately 1 g of fresh plant tissue (corolla excluding the reproductive organs and leaf discs prepared with a 2 cm diameter hole punch) was subjected to two sequential 1-minute washes in 10 mL of n-hexane containing 200 ng of α-amyrin as internal standard. The resulting extracts were vacuum-dried and subsequently saponified in 500 μL of a saponification buffer (20% KOH (w/v) in 50% EtOH (v/v) with 0.5 mg·mL-1 butylated hydroxytoluene) for 2h at 65°C. Subsequently, 100 μL of 10M HCl was added to the aqueous solution to lower the pH to below 2.0, and the mixture was extracted three times with 500 μL of hexane. The resulting extracts were vacuum-dried and subjected to derivatization using a mixture of N-methyl-N-trimethylsilyl-trifluoroacetamide (MSTFA) and N, N-dimethylformamide (DMF) before GC-MS analysis.

For the preparation of pollen and thecal samples, flowers with opened corollas with matured anthers were selected, and anther heads were collected into 5 mL Eppendorf tubes (EP tubes). Following the addition of 1 mL of double-distilled water (ddH2O), samples were vortexed for 30 seconds followed by a 5-minute incubation period. The thecae were then separated into another 5 mL EP tube using a small spatula. After centrifugation at 11,000 g for 15 minutes at 4°C to remove the water layer, two steel beads (2 mm diameter) were added to the tube. The sample was rapidly frozen in liquid nitrogen and homogenized in a ball mill (Genogrinder 2000; SPEX CertiPrep) for 60 seconds at a rate of 1100 strokes per minute.

For triterpene extraction of pollen, thecae and *N. benthamiana* leaves, frozen and ground fresh samples (30 mg for pollen and thecae, and 100 mg for *N. benthamiana* leaves) with 200 ng α-amyrin internal standard were saponified in 300 μL of the saponification buffer described above at 65°C for 2h. Subsequently, 70 μL of 10M HCl was added to the aqueous solution to lower the pH below 2.0, followed by three extractions with 300 μL of hexane. The extracts were then concentrated and derivatized with MSTFA and DMF before GC-MS analysis.

The GC-MS analysis was conducted using the same instrument, columns, temperature programs, and MS settings as in previous studies (Yang et al., 2023).

### Statistical analysis

Statistical analysis using IBM SPSS Statistics 23 (IBM Inc., Chicago, IL, USA). Statistical differences among groups were determined using ANOVA followed by Tukey’s Honestly Significant Difference (HSD) *post hoc* test. The significance of differences between two sample groups was assessed using the student’s t-test. A significance level of *p* ≤ 0.05 was considered statistically significant for all comparisons. The box plots and correlation analysis were conducted using Chiplot (https://www.chiplot.online/).

### Accession numbers

Sequence data from this article can be found in the GenBank/EMBL data libraries under accession numbers NaOSC1 (LOC109226501), NaOSC2 (LOC109226503), CYP716A419 (LOC109234271), CYP716E107 (LOC109209296), CYP716E108 (LOC109209295), CYP716C87 (LOC109239546), CYP716D93 (LOC109205849), CYP716D94 (OIT28874), CYP716H6 (LOC109230529), CYP716A420 (LOC109236645), CYP88B4 (LOC109228169), CYP88C16 (LOC109244115).

## Supplemental Data

**Supplemental Table S1.** Cytochrome P450 enzymes used for phylogenetic tree.

**Supplemental Table S2.** Cytochrome P450 enzymes from *Nicotiana attenuata.*

**Supplemental Table S3.** The level of confidence for β-amyrin derivates identification.

**Supplemental Table S4.** The level of confidence for lupeol and lupanediol derivates identification.

**Supplemental Table S5.** Primer sequences used for the design of constructs for transient expression.

**Supplemental Table S6.** Primer sequences for VIGS.

**Supplemental Table S7.** Primer sequences for qPCR.

**Supplemental Fig. S1.** p450 enzyme from *N. attenuata* plants.

**Supplemental Fig. S2.** Sequence alignment of *N. attenuata* CYP450 candidates with CYP450 enzymes from the triterpenoid biosynthetic pathway.

**Supplemental Fig. S3.** The levels of triterpenes after co-expression of AtLPU1/NaOSC2 with different CYP450s.

**Supplemental Fig. S4** depicts the triterpenoid metabolite profiles of *N. benthamiana* leaves expressing either empty vector (EV) or NaCYP716A419.

**Supplemental Fig. S5.** The levels of triterpenes after co-expression of NaOSC2 and NaCYP716A419 with NaCYP716E107, SlCYP716E26, or NaCYP716C87.

**Supplemental Fig. S6.** The levels of triterpenes after co-expression of AtLPU1 and NaCYP716A419 with NaCYP716E107 or NaCYP716C87.

**Supplemental Fig. S7.** Primary fragmentation pathways of TMS-derivatives of β-amyrin.

**Supplemental Fig. S8.** Primary fragmentation pathways of TMS-derivatives of erythrodiol.

**Supplemental Fig. S9.** Primary fragmentation pathways of TMS-derivatives of oleanolic aldehyde.

**Supplemental Fig. S10.** Primary fragmentation pathways of TMS-derivatives of oleanolic acid.

**Supplemental Fig. S11.** Primary fragmentation pathways of TMS-derivatives of maslinic acid.

**Supplemental Fig. S12.** Primary fragmentation pathways of TMS-derivatives of lupeol.

**Supplemental Fig. S13.** Primary fragmentation pathways of TMS-derivatives of lupanediol.

**Supplemental Fig. S14.** Primary fragmentation pathways of TMS-derivatives of betulin.

**Supplemental Fig. S15.** Primary fragmentation pathways of TMS-derivatives of betulinic acid.

**Supplemental Fig. S16.** Alignment of protein sequences of NaCYP716E107, SlCYP716E26, and CaCYP716E41.

**Supplemental Fig. S17.** The levels of triterpenes after expressed NaCYP716E107 or SlCYP716E26.

**Supplemental Fig. S18.** Silencing efficiency of 4 triterpene biosynthesis genes in VIGS plants.

**Supplemental Fig. S19.** 2,3,5-triphenyltetrazolium chloride (TTC) staining of pollen from EV (empty vector), NaOSC1/2-VIGS, and CYP716A419-VIGS plants.

**Supplemental Fig. S20.** GC-MS chromatograms of extracts of *N. attenuata* roots with Phytohormone treatments.

**Supplemental Fig. S21.** Products of NaCYP716A419, NaCYP716C87, and NaCYP716E107 in *N. attenuata* roots (infected with arbuscular mycorrhizal fungi (AMF) for 5 weeks).

**Supplemental Fig. S22.** GC-MS chromatograms of reaction products of NaOSC2/CYP716A419 co-expressed with other CYP450 candidates, aligned with standards.

**Supplemental Fig. S23.** GC-MS chromatograms of reaction products of NaOSC2/CYP716A419/CYP716C87 or NaOSC2/CYP716A419/CYP716E107 co-expressed with other CYP450 candidates in *N. benthamiana*, aligned with authentic standards.

## Acknowledgments

We acknowledge Dr. Xincong Jiang for photographing the pollen. We acknowledge Prof. David R. Nelson for his contributions in naming candidate CYP450 enzymes in the manuscript.

## Author contributions

CY conceived the study, conducted the experiments, analyzed the data. ITB and SEO supervised the project. CY and ITB wrote the manuscript. RH provided valuable technical support for metabolite analysis. All the authors commented on the manuscript.

## Data availability

All data are incorporated into this article and its online supplementary material.

